# Microbial eco-evolutionary dynamics of decomposition and dormancy

**DOI:** 10.64898/2026.09.06.749690

**Authors:** Agathe Chave-Lucas, Régis Ferrière

## Abstract

Soil microorganisms regulate a major component of the terrestrial carbon cycle, yet predictions of soil carbon stocks and fluxes often neglect microbial life-history adaptation. To fill this gap, we develop a spatially and stage-structured eco-evolutionary model in which active and dormant microbes move between favorable microsites and an unfavorable bulk soil matrix; decompose organic carbon through costly exoenzyme production; and evolve both exoenzyme investment and entry into dormancy. The model shows that dormancy expands the ecological conditions under which microbial populations persist and has a non-monotonic effect on soil carbon stocks. Adaptive dormancy is shaped by opposing selection in microsites, where inactivity carries an opportunity cost, and in the matrix, where dormancy protects cells from mortality. When dormancy and exoenzyme production jointly evolve, the traits may increase together under high microbial mobility, but often evolve in opposite directions because both carry survival benefits in the matrix. These eco-evolutionary feedbacks can either amplify or attenuate soil carbon fluxes to the atmosphere, depending on soil structure, microbial movement, and dormancy costs. Our results suggest that incorporating microbial life-history evolution into soil carbon models is essential for predicting soil carbon feedbacks to climate.

## Introduction

Soils are central to Earth’s carbon cycle, as the ecological and biogeochemical processes they sustain regulate the storage and release of carbon. They store carbon in both organic and in-organic forms, and the total soil carbon reservoir is estimated to be approximately twice the size of the atmospheric carbon pool (Scharlemann et al. 2014; Ontl and Schulte 2012). As a result, even modest changes in soil carbon storage can translate into substantial changes in atmospheric carbon concentrations, with important consequences for future climate.

Soil organic carbon originates largely from plant and microbial residues that are progressively transformed by soil organisms (Singh et al. 2010). Soil fauna such as earthworms, fragment litter and mix organic residues with mineral particles, increasing the surface area available for microbial attack. Microbial communities then enzymatically depolymerize and metabolize this material, releasing carbon dioxide and other compounds such as methane as by-products of respiration and anaerobic metabolism. Through their control of organic matter decomposition, soil microorganisms therefore exert a major influence on greenhouse gas fluxes from soils to the atmosphere (Raich and Schlesinger 1992, Schimel 1995).

Microbial control of decomposition is not fixed. Owing to their enormous population sizes, short generation times, and substantial genetic diversity, microbial communities can adapt rapidly to changing environmental conditions (Padfield et al. 2015, Schaum et al. 2017). Yet the mechanisms by which adaptive microbial traits evolve, and the extent to which this evolution feeds back on ecosystem processes such as organic matter decomposition and CO_2_ respiration, remain insufficiently understood (Classen et al. 2015, Abs, Leman, and Ferrière 2020, Abs, Chase, et al. 2024, Greenblum 2024).

One feature that is likely to be especially important, but remains incompletely integrated into current frameworks of microbial adaptation, is the spatial organization of microbial communities (Nunan, Schmidt, and Raynaud 2020). Soils are highly heterogenous three-dimensional systems composed of solid particles interspersed with water- and air-filled pore networks. Within this complex architecture, resource availability, water potential, oxygen, and cell density can vary over micrometers to millimeters, and microbial activity is concentrated in small, spatially clustered communities associated with soil aggregates (Paul and Clark 1996, Ettema and Wardle 2002, Young and Crawford 2004, Nunan 2017).

Soil aggregates form a patchwork of favorable microsite embedded in the bulk soil, where environmental conditions are generally less suitable for microbial activity and cells may remain physically isolated for extended periods. Microorganisms tend to remain within microsites (Wilpiszeski et al. 2019), but they can also move between microsites and the bulk soil, and through the bulk soil itself. Such movement is thought to occur predominantly through passive transport processes (Abu-Ashour et al. 1994), whereas active motility is strongly constrained outside the wetter range of matrix potentials because of its high energetic cost (Dechesne et al. 2010). Together, these features raise fundamental questions about how soil structure and microbial movement between microsites affect ecosystem functions such as decomposition and respiration, and how these relationships vary across macroscopic soil conditions.

Despite this spatial complexity, many theoretical models represent soils as a single well-mixed compartment in which substrates and microbes diffuse freely (Abs and Ferrière 2020; Chandel, Jiang, and Luo 2023). This simplification can provide a useful first approximation, but it may be misleading when the traits under study experience different selection pressures in different microenvironments (Özkaya et al. 2017; Bonner et al. 2022; Chave-Lucas and Ferrière n.d.). Exoenzyme production is one such trait. Exoenzymes are costly extracellular enzymes that depolymerize soil organic matter into dissolved organic matter that microbes can uptake and assimilate. They are therefore crucial to soil carbon decomposition and, more broadly, to soil carbon cycling. Because extracellular exoenzymes and the monomers they generate can diffuse away from their producers, exoenzyme production has features of a public good. This entails that cells that produce less or no exoenzymes can benefit from enzymes produced by others while avoiding the cost of enzyme synthesis. In hydrated microsites, diffusion and local sharing can therefore favor reduced investment in exoenzyme production. In contrast, in the bulk soil, where cells are much more isolated and diffusion is limited, producers may retain a larger fraction of the benefits of their own exoenzymes, favoring higher investment in exoenzyme production (Chave-Lucas and Ferrière n.d.; Allison 2005; Sinsabaugh et al. 2008; Burns et al. 2013).

Microbial dormancy may establish an analogous temporal contrast in selection (Zhang et al. 2022). Dormancy in microbes is a reversible state of strongly reduced growth and metabolism, reduced mortality, and increased resistance to environmental stress. Dormant cells can persist for months to years, and in some cases much longer (Lennon and Jones 2011; Bradley 2025). The ability to enter dormancy can be costly because it requires physical machinery for sensing environmental conditions, switching metabolic state, and maintaining cellular integrity during prolonged inactivity. In unfavorable bulk soil, dormancy can be favored because it allows cells to reduce mortality or maintenance costs while waiting for improved conditions. In resource-rich, hydrated microsites, by contrast, dormancy entails an opportunity cost because active cells can grow and divide; selection may therefore favor lower investment in dormancy or more rapid awakening where microbes are expected to switch to dormancy, and under negative selection in microsites, where microbes should awaken (Zhang et al. 2022; Lennon and Jones 2011; Blagodatskaya and Kuzyakov 2013; Joergensen and Wichern 2018).

Many soil models, including models that explicitly represent microbial biomass and spatial heterogeneity, omit dormancy because dormant microbes are often assumed to contribute little to carbon cycling. Dormant microbes consume few resources, and their carbon emissions are usually considered negligible relative to those of active cells. However, recent studies suggest that incorporating dormancy into soil carbon models can improve predictions of microbial biomass and soil carbon decomposition (He et al. 2015). This is because dormant cells can dominate microbial biomass: in some soils, as much as 80 or 90% of the microbial community may be dormant at any given time. Moreover, the active-to-dormant ratio and the investment required to enter, maintain, and exit dormancy are expected to change with environmental warming (McDonald et al. 2024), with potentially large consequences for carbon fluxes to the atmosphere.

Here, we develop a spatially heterogeneous mathematical model that couples soil carbon decomposition, microbial movement, and dormancy in a patchwork of favorable microsites embedded in unfavorable bulk soil. The eco-evolutionary model captures local feedbacks between microbial populations and their environment and predicts how these local dynamics scale up to affect soil-atmosphere carbon exchange. We focus on two evolving traits: investment in exoenzyme production, which directly controls access to soil organic carbon, and the propensity to enter dormancy, which controls persistence under unfavorable conditions. We evaluate the evolution of these traits and their joint evolution and ask how adaptive microbial responses alter the soil-atmosphere carbon feedback under environmental change.

## Methods

Soil microbial dynamics are heterogeneous in both space and time. Spatially, microbial growth and decomposition are concentrated in small soil aggregates, or microsites, embedded in a less favorable bulk soil matrix. Temporally, microbial cells may alternate between active and dormant physiological states, with dormancy allowing cells to persist through adverse environmental conditions. We therefore develop a spatially and stage-structured model that tracks active and dormant microbes in microsites and in the bulk soil matrix, together with soil organic carbon, dissolved organic carbon, and exoenzymes.

### Ecological model

To describe the dynamics of active and dormant microbes and their effects on soil carbon stocks, we extend a CDMZ-type ecological model (Abs, Saleska, et al. 2025; Chave-Lucas and Ferrière n.d.). The model explicitly represents active and dormant microbial biomass in microsites (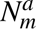 and 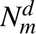 ) and in the bulk soil matrix (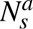 and 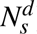), where subscripts *m* and *s* denote microsites and bulk soil, respectively, and superscripts *a* and *d* denote active and dormant individuals. The model also tracks the concentration of exoenzymes produced by active microorganisms in microsites (*Z_m_*), soil organic carbon in microsites (*C_m_*), dissolved organic carbon in microsites (*D_m_*), and non-dissolved organic carbon in the bulk soil matrix (*C*). In microsites, exoenzymes depolymerize soil organic carbon into dissolved organic carbon, which active microorganisms can assimilate. In the bulk soil matrix, active microbes do not grow but may die, enter dormancy, awaken, or move between the matrix and microsites.

The ecological dynamics are governed by a system of eight ordinary differential equations

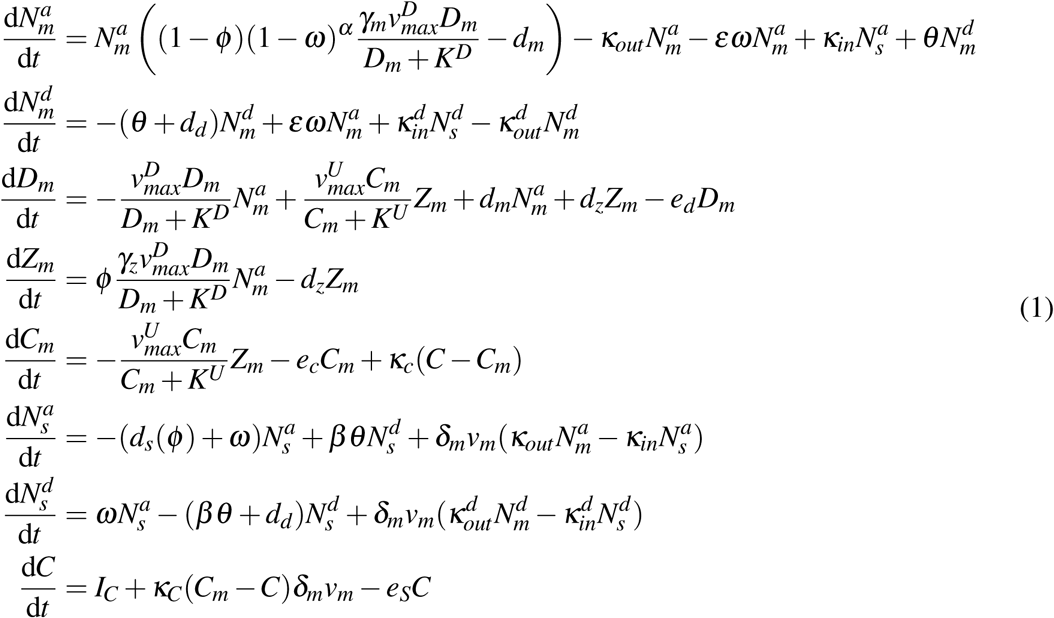

Following Chave-Lucas and Ferrière n.d., *γ_m_* is the microbial growth efficiency, *γ_z_* is the enzyme production efficiency, *d_m_* is the death rate of active microbes, *d_d_* is the death rate of dormant microbes, and *d_z_*is the denaturation rate. Microbial uptake of dissolved organic carbon, exoenzyme production, The growth of microorganisms in microsites, the production of exoenzymes, and enzymatic production of dissolved organic carbon are described described by Michaelis-Menten functions, in which *v_max_* and *K* denote the maximum reaction rate and the half-saturation constant, respectively. Parameters *e_c_*, *e_d_*, and *e_s_* denote the loss rates of soil organic carbon in microsites, dissolved organic carbon in microsites, and soil organic matter in the bulk soil matrix, respectively; *I_C_* is the input rate of soil organic matter to the bulk soil matrix.

The trait *φ* denotes the fraction of assimilated dissolved organic carbon that is invested in exoenzyme production. Each active microbial cell therefore allocates a fraction *φ* of its assimilated carbon into exoenzyme production, and the remaining fraction, (1 − *φ* ), to growth, assuming that the minimum carbon requirement for cell maintenance is always met. The trait *ω* denotes the rate of entry into dormancy in the bulk soil, while *θ* denotes the rate of exit from dormancy in microsites. In the equations above, entry into dormancy in microsites is scaled by *εω*, and exit from dormancy is scaled by *βθ* in the bulk soil.

As in Tang and Riley’s model (Tang and Riley 2019), parameters *δ_m_*and *v_m_* represent the density and volume of microsites. The parameter *κ_c_* is the conductance of organic matter between microsites and the bulk soil matrix. The parameters *κ_in_* and *κ_out_* are conductance rates for active microbes, with *κ_in_*denoting movement from the bulk soil matrix into microsites and *κ_out_* denoting movement out of microsites. Dormant microbes move analogously, with conductance rates 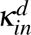 and 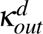. Because microbes inside microsites are assumed to adhere to each other, we generally assume *κ_in_* to be larger than *κ_out_*; similarly, 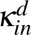 is assumed to be larger than 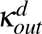).

Both focal traits, exoenzyme investment (*φ* ) and entry into dormancy (*ω*), carry costs for active microbial growth. These costs are represented by the multiplicative factor (1 − *φ* )(1 − *ω*)*^α^* in the microsite growth term, where *α* controls the shape and strength of the cost of dormancy investment. Environmental conditions, especially temperature and soil moisture, are expected to affect both entry into and exit from dormancy. More favorable conditions should reduce entry into dormancy and increase exit from dormancy, whereas unfavorable conditions should have the opposite effect.

In microsites, where water and nutrients are more readily available, microbes grow and produce exoenzymes. In the bulk soil matrix, by contrast, active microbes are assumed to grow less than they die, as in Chave-Lucas and Ferrière n.d. The mortality rate of active microbes in the bulk soil matrix depends on exoenzyme investment, *φ* , and is thus denoted by *d_s_*(*φ* ). We use the same functional form as in Chave-Lucas and Ferrière n.d.

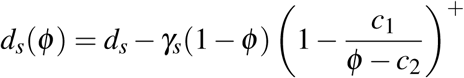

where *d_s_* is the basal death rate of microbes in the bulk soil, *γ_s_* controls the maximum reduction in mortality, *c*_1_ and *c*_2_ shape the dependence of mortality on *φ* , and (*x*)^+^ = max(*x,* 0). Parameter *γ_s_* is assumed to always be strictly smaller than *d_s_*, ensuring that *d_s_*(*φ* ) is always negative. This formulation captures the assumption that exoenzyme investment can reduce mortality in the bulk soil matrix, for example by increasing access to resources (dissolved organic carbon) and thereby increasing the chance that a cell survives until it reaches a favorable microsite.

Microbes can enter and exit dormancy in both microsites and the bulk soil matrix. We assume that entry into dormancy is more likely in the bulk soil, where active cell mortality is high, whereas exit from dormancy is more likely in microsites, where growth is possible. Following J. A. J. Metz, Klinkhamer, and Jong 2009, exit from dormancy in the bulk soil is proportional to exit from dormancy in microsites through the parameter *β* , and entry into dormancy in microsites is proportional to entry into dormancy in the bulk soil through the parameter *ε*. We therefore use *ε <* 1 and *β <* 1 as default assumptions, while also exploring other parameter ranges numerically.

### Invasion fitness

Because the ecological model is structured both spatially and physiologically, standard expressions for invasion fitness in homogeneous populations (Parvinen and Seppänen 2016) cannot be applied directly. Instead, the calculation must account for the differential growth of mutant biomass in microsites and in the bulk soil, and for both active and dormant states.

We first consider the evolution of the dormancy entry rate, *ω*, while holding the exoenzyme investment, *φ* , fixed. A rare mutant with trait *ω_mut_* arises in a resident population with trait *ω* at ecological equilibrium. Because the mutant is initially rare, the mutant growth equations are linear and are given by

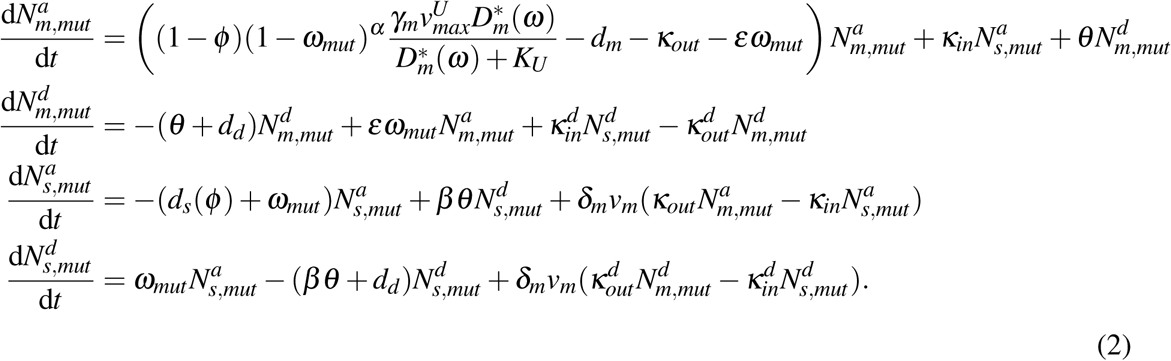

Here, 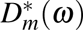 is the equilibrium concentration of dissolved organic carbon in microsites generated by the resident population with trait *ω*, obtained from Eq. 1. Building on adaptive dynamics theory J. Metz, Nisbet, and Geritz 1992 and on the approach in Chave-Lucas and Ferrière n.d., invasion fitness for the evolution of *ω* is the dominant eigenvalue of the linear mutant system, Eq. 2.

We denote this invasion fitness by *ρ*(*ω, ω_mut_*). It is the largest eigenvalue of the matrix

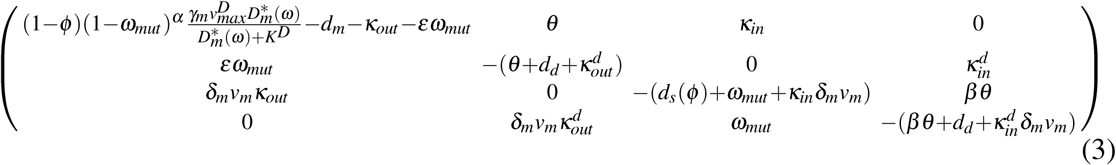

Evolutionary singularities for *ω*, denoted by *ω*\*, are candidate attractors or repellors of the adaptive dynamics and are obtained as zeros of the selection gradient:

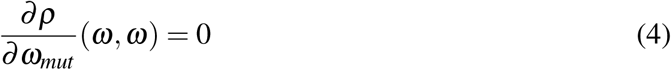

We next consider the joint evolution of entry into dormancy, *ω*, and exoenzyme investment, *φ* . The invasion fitness *f* (*ω, φ, ω_mut_, φ_mut_*) of a rare mutant with traits (*ω_mut_, φ_mut_*) in a resident population with traits (*ω, φ* ), is again the dominant eigenvalue of corresponding linear mutant system. In this case, it is the largest eigenvalue of the matrix

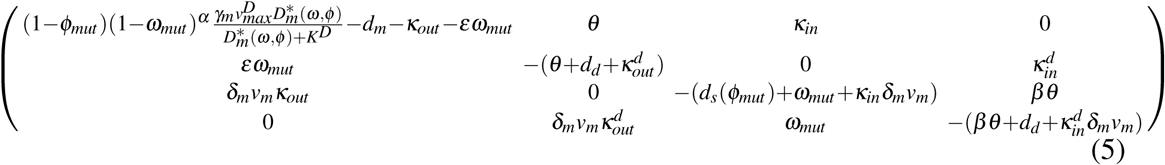

Joint singularities, denoted by (*ω*\**, φ* *), are candidate attractors or repellors of the two-dimensional adaptive dynamics and satisfy

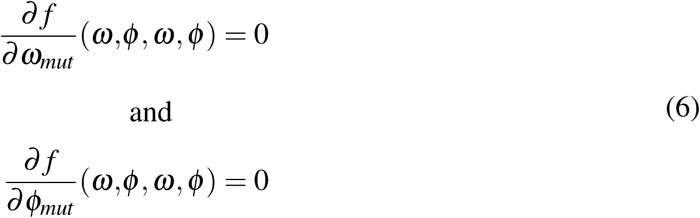

### Default parameters

Unless otherwise noted, default values for the microbial, enzymatic and carbon-cycling parameters (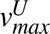, 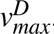, *K^U^* , *K^D^*, *γ_m_*, *γ_z_*, *d_z_*, *d_m_*, *e_d_*, *e_s_*, *e_c_* and *I_C_*) follow Abs, Saleska, et al. 2025. Approximate values for the microsite volume fraction, *δ_m_v_m_*, and for carbon conductance, *κ_c_*, follow Tang and Riley 2019. Values of *κ_in_*, *κ_out_*, *d_s_*, *c*_1_, *c*_2_ and *γ_s_* follow Chave-Lucas and Ferrière n.d. We assume 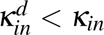 and 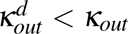 because dormant microbes are expected to be less mobile than active microbes, and we assume a very small dormant-microbe death rate, *d_d_* . Default values of *ε* and *β* are less than one, reflecting lower dormancy entry in microsites than in the bulk soil matrix and lower dormancy exit in the bulk soil matrix than in microsites. We center *α* near one and *θ* near 10^−2^, while exploring broader ranges of the parameters in numerical analyses.

## Results

To understand how dormancy and spatial population structure shape eco-evolutionary dynamics, we first identify the conditions under which the ecological system is viable and quantify the ecological effect of dormancy on soil carbon stocks. We then analyze how the evolutionarily adapted rate of entry into dormancy changes with soil and microbial parameters. Finally, we examine the joint evolution of dormancy and exoenzyme investment across the same parameter gradients.

### Ecological viability and extinction

We first determine the combinations of dormancy entry, *ω*, and exoenzyme investment, *φ* , for which the microbial system persists. Under some conditions, especially when both *ω* and *φ* are low, the microbial population becomes extinct. Extinction occurs either because active cells experience excessive mortality while moving through the bulk soil matrix or because insufficient exoenzyme production prevents the conversion of soil organic carbon into dissolved organic carbon, thereby limiting microbial growth.

A similar analysis was performed by Chave-Lucas and Ferrière n.d. in the absence of dormancy. Adding dormancy broadens the viability domain of the system. In particular, without dormancy, high microbial mobility can drive the entire population extinct, whereas survival remains possible at higher mobility when cells can enter dormancy. For some parameter ranges, even very low rates of entry into dormancy are sufficient to ensure persistence (Fig. 1A,B), whereas all microbes go extinct in the corresponding system without dormancy. In other parameter ranges, including those explored by Chave-Lucas and Ferrière n.d., dormancy is not required for persistence, except, potentially, when exoenzyme investment is very high (Fig. 1C).

**Figure 1.**
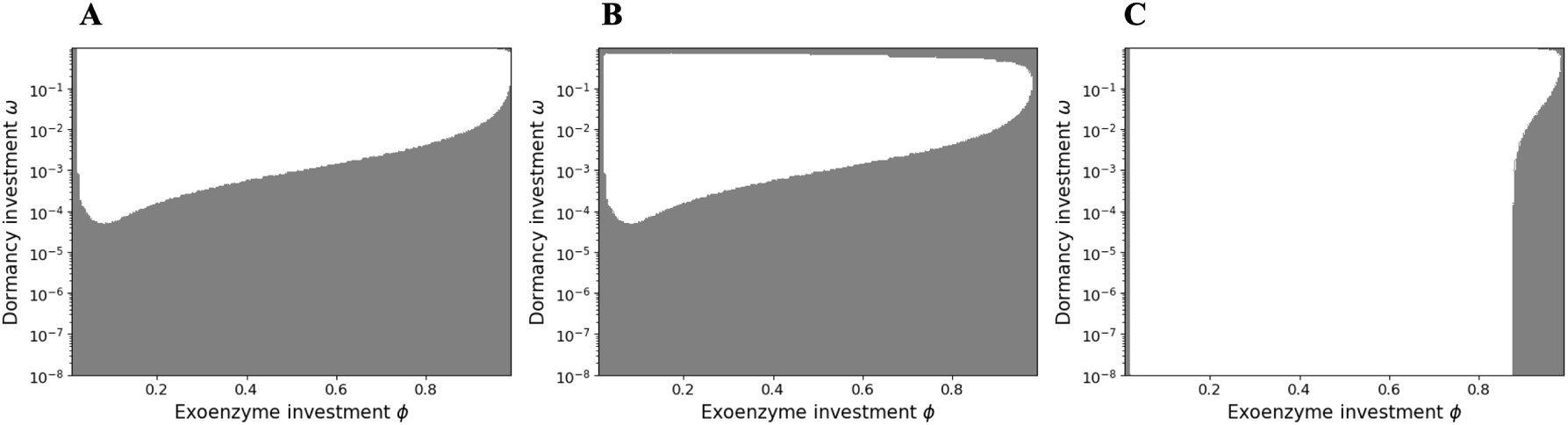
Dormancy, exoenzyme investment, and ecological viability. White regions indicate parameter combinations for which the microbial population persists; gray regions indicate extinction. In all panels, *θ* = 0.4 and *δ_m_v_m_*= 3.9 × 10^−4^. Unless otherwise specified, all other parameters are given in Table 1. **A** Extinction occurs when both exoenzyme investment and dormancy entry are low. Here, *α* = 1, 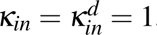, and 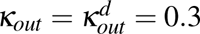. **B** Same microbial parameters as in **A**, but with *α* = 5. In this case, the microbial population also becomes extinct when dormancy entry and exoenzyme investment are high. **C** *κ_in_* = 1, 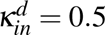, *κ_out_* = 10^−2^, and 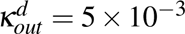 and *d_s_* = 3 × 10^−2^. Except at very high values of both dormancy entry and exoenzyme investment, the microbial population persists whether or not dormancy occurs.

**Table 1.** Parameter list and default values.

| Parameter | Description | Unit | Standard value |
| --- | --- | --- | --- |
| $\gamma_m$ | microbial growth efficiency | | 0.2 |
| $\gamma_s$ | microbial bulk soil growth term | $\text{h}^{-1}$ | $2 \times 10^{-4}$ |
| $\gamma_z$ | enzyme production efficiency | | 0.2 |
| $d_m$ | active microbial microsite death rate | $\text{h}^{-1}$ | $1.5 \times 10^{-4}$ |
| $d_s$ | active microbial bulk soil death rate | $\text{h}^{-1}$ | $3 \times 10^{-4}$ |
| $d_d$ | dormant microbial death rate | $\text{h}^{-1}$ | $10^{-6}$ |
| $d_z$ | enzyme degradation rate | $\text{h}^{-1}$ | $2 \times 10^{-3}$ |
| $v_{max}^U$ | maximum microbe reaction rate | $\text{h}^{-1}$ | $4 \times 10^{-1}$ |
| $v_{max}^D$ | maximum enzyme reaction rate | $\text{h}^{-1}$ | $4 \times 10^{-1}$ |
| $K^U$ | microbe half-saturation constant | $\text{mg C cm}^{-3}$ | $10^{-1}$ |
| $K^D$ | enzyme half-saturation constant | $\text{mg C cm}^{-3}$ | 50 |
| $e_d$ | decomposed carbon leaching rate | $\text{h}^{-1}$ | $10^{-2}$ |
| $e_c$ | microsite carbon leaching rate | $\text{h}^{-1}$ | $10^{-6}$ |
| $e_s$ | bulk soil carbon leaching rate | $\text{h}^{-1}$ | $10^{-6}$ |
| $I_C$ | organic carbon input | $\text{mg C cm}^{-3} \text{ h}^{-1}$ | $10^{-3}$ |
| $\delta_m v_m$ | fraction of soil occupied by microsites | | $3.9 \times 10^{-3}$ |
| $\kappa_c$ | carbon soil conductance | $\text{h}^{-1}$ | 10 |
| $\kappa_{in}$ | inward active microbial flux rate | $\text{h}^{-1}$ | 1 |
| $\kappa_{out}$ | outward active microbial flux rate | $\text{h}^{-1}$ | $10^{-2}$ |
| $\kappa_{in}^d$ | inward dormant microbial flux rate | $\text{h}^{-1}$ | 0.5 |
| $\kappa_{out}^d$ | outward dormant microbial flux rate | $\text{h}^{-1}$ | $5 \times 10^{-3}$ |
| $\gamma_s$ | bulk soil uptake rate | $\text{h}^{-1}$ | $2 \times 10^{-4}$ |
| $c_1$ | bulk soil growth constant | | $10^{-2}$ |
| $c_2$ | bulk soil growth constant | | $10^{-2}$ |
| $\epsilon$ | microsite dormancy constant | | $10^{-1}$ |
| $\theta$ | microsite awakening rate | $\text{h}^{-1}$ | $10^{-2}$ |
| $\beta$ | bulk soil dormancy constant | | $10^{-6}$ |
| $\alpha$ | dormancy cost constant | | 1 |

### Effect of dormancy on carbon stocks

Dormancy entry not only affects microbial persistence but also has a nonlinear effect on soil carbon stocks (Fig. 2A). As *ω* initially increases, soil carbon stocks decline *ω* reaches a threshold value (approximately 0.1 in our numerical example), after which they rise again. At low values of *ω*, increased dormancy allows more microbes to survive adverse conditions in the bulk soil, thereby increasing decomposition and lowering soil carbon stocks. At higher values of *ω*, however, a larger fraction of the population remains dormant. Because dormant cells do not contribute to decomposition, carbon stocks then increase.

**Figure 2.**
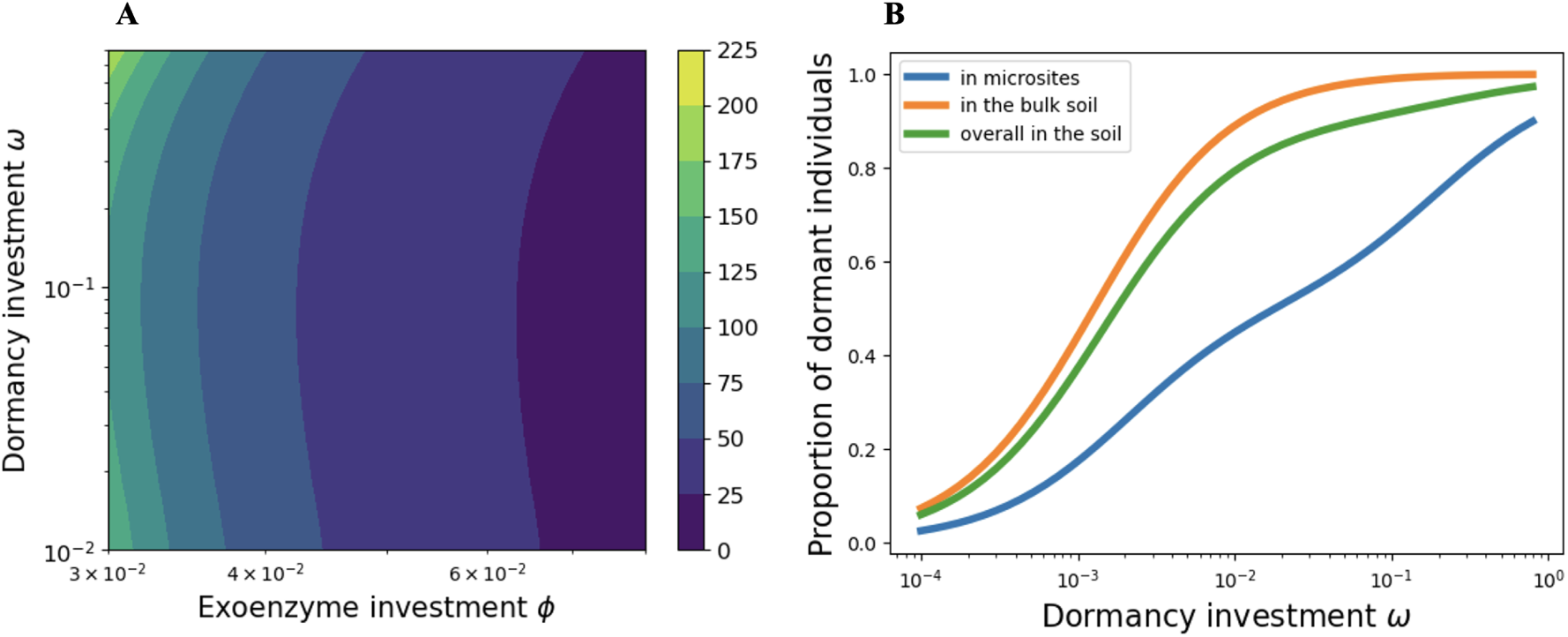
Effects of dormancy and soil carbon and microbial activity. **(A)** Equilibrium soil carbon stock in microsites, 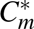 (mg C cm^−3^), as a function of dormancy entry, *ω*, and exoenzyme investment, *φ* . **(B)** Fraction of dormant microbes as a function of dormancy entry, *ω*, for *φ* = 0.1. See Table 1 for other parameter values.

Empirical studies suggest that dormant microbes can represent 80–95% of total microbial biomass Zhao et al. 2026. For the parameter values used in Fig. 2A, this empirical range corresponds approximately to values of *ω* between 10^−2^ and 10^−1^ (Fig. 2B). This is also the range over which carbon stocks initially decline in Fig. 2A, suggesting that variation in dormancy entry within empirically plausible bounds could substantially affect soil carbon storage.

### Evolution of entry into dormancy

We next analyze how the dormancy-entry trait, *ω*, evolves as soil and microbial parameters vary. For parameters near the default set (Table 1), the system generally evolves toward a viable evolutionary singularity, denoted by *ω*\*, which represents the adapted rate of entry into dormancy. Across the parameter gradients considered here, *ω*\* ranges from approximately 0.04 and 0.15 (Fig. 3). This range overlaps with the interval over which dormancy has strong effects on soil carbon stocks (Fig. 2), and most of the variation in *ω*\* occurs at relatively low exoenzyme investment (0.03 *< φ <* 0.05).

**Figure 3.**
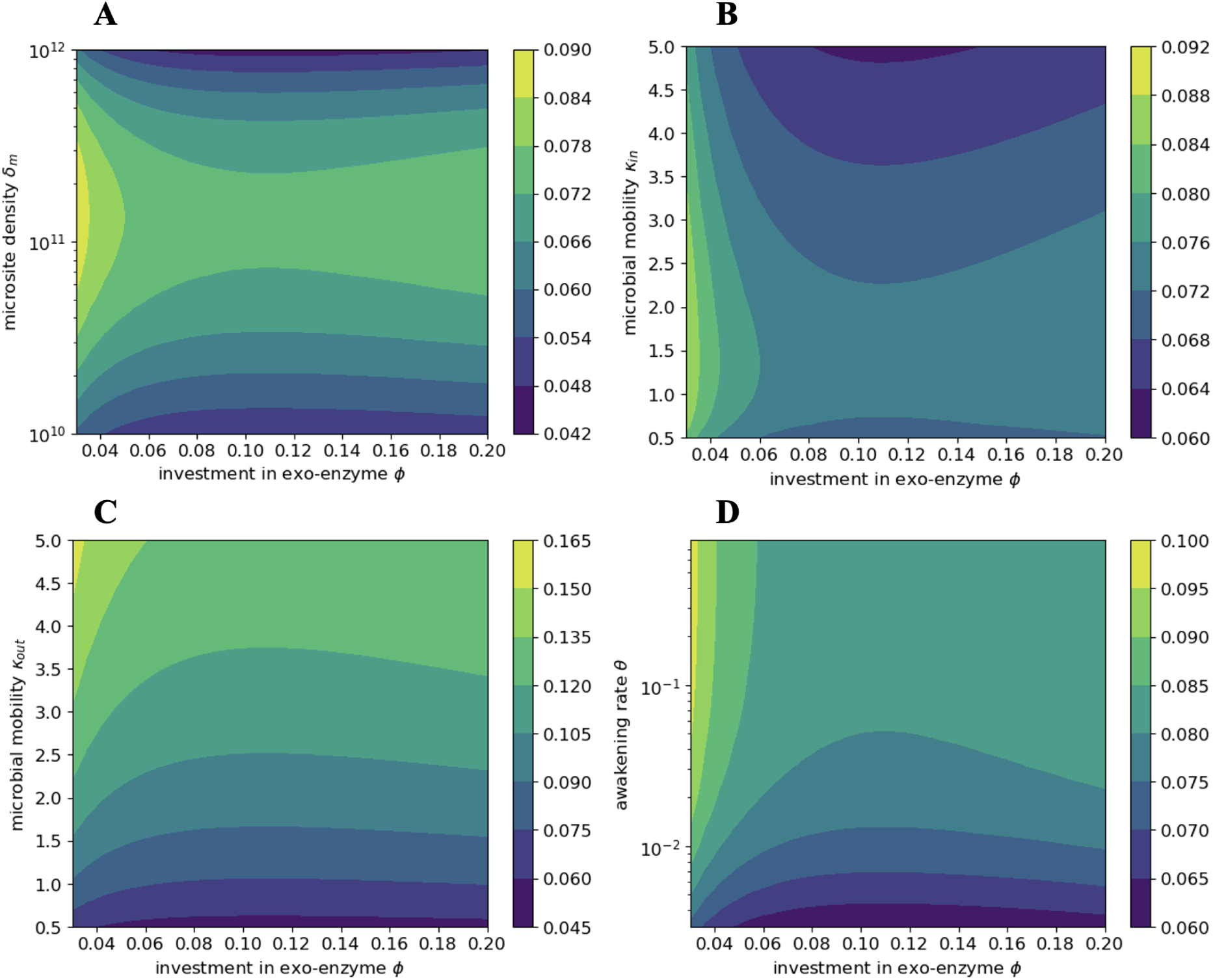
Evolutionarily adapted dormancy entry. Adapted investment in dormancy, *ω*\*, as a function of exoenzyme investment, *φ* soil and **(A)** microsite density *δ_m_*(m^−3^); **(B)** microbial mobility into microsites, *κ_in_*, with *κ_out_* fixed; **(C)** total microbial mobility, with the ratio *κ_in_/κ_out_* held constant; and **(D)** awakening rate, *θ* . See Table 1 for other parameter values.

The evolutionary singularity *ω*\* varies non-monotonically with exoenzyme investment *φ* (Fig. 3). As *φ* increases up to approximately 0.1, *ω*\* declines, indicating that dormancy becomes less advantageous when cells invest more in exoenzyme production and are therefore better able to survive in the bulk soil. At higher values of *φ* , however, *ω*\* increases again. In this range, the cost of exoenzyme production becomes sufficiently high that increased dormancy is favored as a way to offset the reduction in active growth driven by the exoenzyme production cost.

Consistent with Chave-Lucas and Ferrière n.d., *ω*\* reaches its highest values at intermediate microsite density, *δ_m_*, around 10^11^ m^−3^ under the parameter values considered here (Fig. 3A).

When microsite density is very low, microbes spend long periods in the bulk soil and often die before reaching another favorable patch, regardless of their dormant strategy; selection for dormancy entry is therefore weak. For microsite densities above 10^11^, by contrast, cells return to favorable microsites more rapidly, and frequent dormancy slows growth more that it improves survival. Accordingly, *ω*\* declines at high microsite density. This intermediate-microsite density maximum depends strongly on the death rate of dormant microbes and disappears when *d_d_ <* 10^−8^. Under extremely low dormant-cell mortality, *ω*\* always decreases monotonically with microsite density (Fig. S1).

Microbial mobility also strongly affects *ω*\*, as in the model of Chave-Lucas and Ferrière n.d. When total microbial mobility increases while the ratio *κ_in_/κ_out_* remains constant, the singularity *ω*\* increases, since microbes will tend to spend more time in the bulk soil where they die more (Fig. 3C). A different pattern emerges when *κ_out_* is fixed and only *κ_in_* varies. In this case, increasing *κ_in_* generally reduces *ω*\* because cells in the bulk soil locate microsites more rapidly, reducing the benefit of dormancy (Fig. 3B).

At low values of *κ_in_* or low microsite density, *δ_m_*, however, *ω*\* can increase with microbial movement into microsites (Fig. 3B and Fig. S2). This pattern reflects a trade-off between mortality in the bulk soil and the time required to encounter a new microsite. When movement is very limited, increasing *κ_in_* reduces the fraction of dormant cells in both microsites and the bulk soil, exposing more active individuals to mortality. In this range, higher dormancy entry compensates the mortality cost and therefore is selected. Once *κ_in_*and microsite density are sufficiently high, the time spent in the bulk soil becomes short enough that mortality risk declines, and lower values of *ω*\* are favored.

As expected, the singularity *ω*\* is also highly sensitive to parameters that directly govern individual dormancy dynamics (Fig. 3D and Fig. S3). In particular, *ω*\* increases with the awakening rate, *θ* . When microbes awaken faster, especially outside microsites, selection favors a higher rate of reentry into dormancy.

### Joint evolution of exoenzyme investment and dormancy

We finally examine the joint evolution of dormancy entry, *ω*, and exoenzyme investment, *φ* , across gradients in microbial and soil parameters. Depending on the parameter varied, the two evolutionary singularities, *ω*\* and *φ* *, can shift in the same direction (Fig. 4B) or in opposite directions (Fig. 4A, 4E, 4F). Across these gradients, *ω*\* remains approximately between 0.04 and 0.15, whereas *φ* * is concentrated between 0.03 and 0.05. This range of *φ* * is narrower than that obtained without dormancy under comparable parameters in Chave-Lucas and Ferrière n.d.’s study. It also coincides with the values of exoenzyme investment for which soil carbon stocks are most sensitive to changes in dormancy (Fig. 2A).

**Figure 4.**
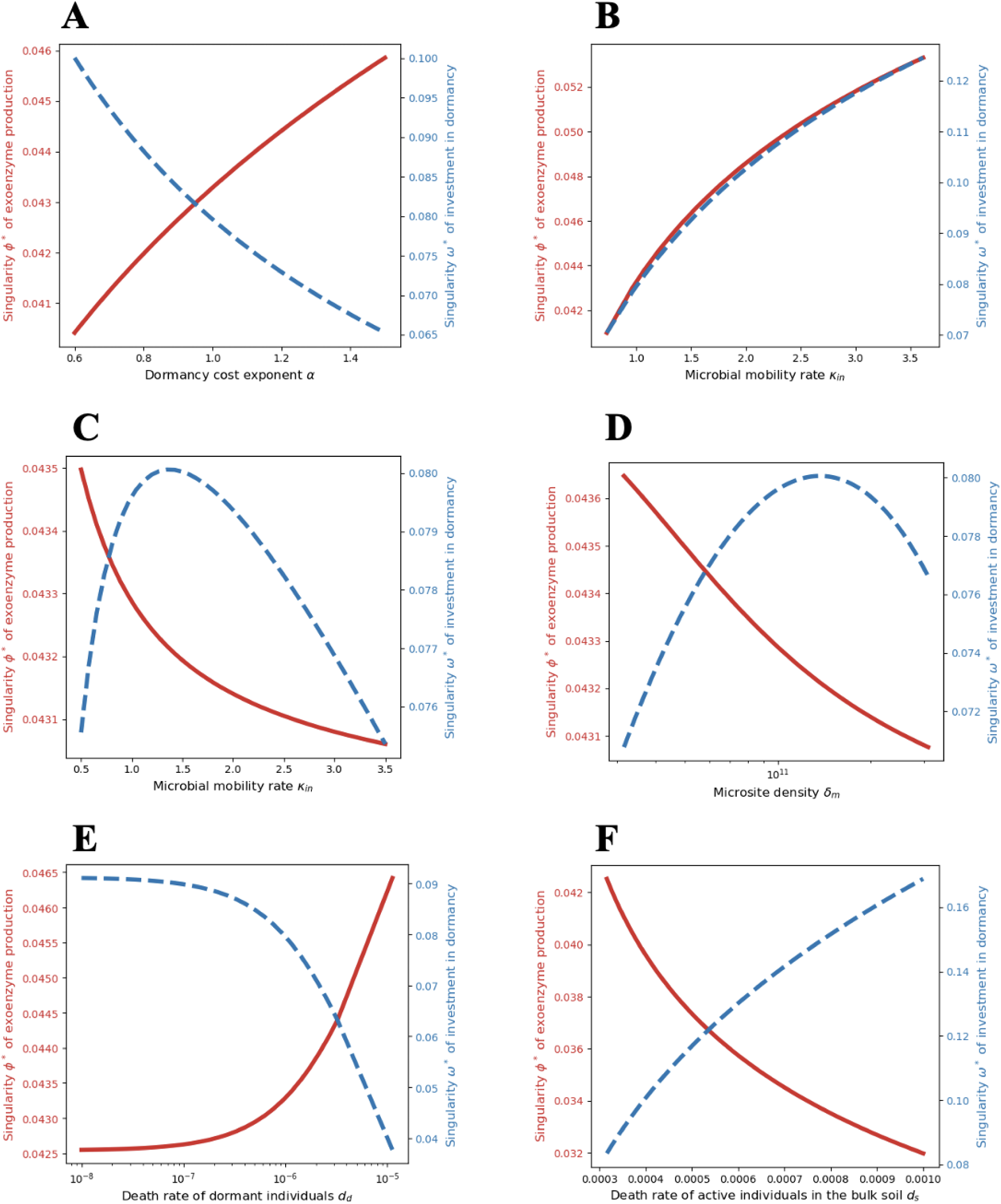
Joint evolution of dormancy and exoenzyme investment. Evolutionarily adapted dormancy entry, *ω*\*, and exoenzyme investment, *φ* *, as functions of **(A)** dormancy-cost exponent, *α*; **(B)** total microbial mobility, with the ratio *κ_in_/κ_out_* held constant; **(C)** microbial mobility into microsites, *κ_in_*, with *κ_out_* held constant; **(D)** microsite density, *δ_m_*; **(E)** death rate of dormant individuals, *d_d_*; and **(F)** death rate of active individuals in the bulk soil, *d_s_*. See Table 1 for values of parameters kept constant.

When the cost of dormancy increases, the two traits evolve in opposite directions (Fig. 4A). As the exponent *α* increases, dormancy becomes more costly, and the adapted rate of dormancy entry, *ω*\*, declines. Lower dormancy entry, however, increases exposure to mortality in the bulk soil. As a result, selection favors higher exoenzyme investment, *φ* *, which improves survival outside microsites.

When the ratio *κ_in_/κ_out_* is held constant, increasing *κ_in_* also increases *κ_out_* and therefore increases overall microbial mobility (Fig. 4B). In this case, cells leave microsites more frequently and are more often exposed to the adverse conditions of the bulk soil. Both dormancy and exoenzyme production therefore increase. By contrast, when *κ_out_* is fixed and only *κ_in_* increases, cells can find new microsites more rapidly without leaving them more often (Fig. 4C). As in Fig. 3B, *ω*\* first increases and then decreases, whereas *φ* * decreases monotonically. The range of variation in *φ* * is small relative to that of *ω*\*.

A similar contrast occurs along the microsite-density gradient (Fig. 4D). The adapted dormancy entry reaches a maximum at intermediate microsite density, as in Fig. 3A, whereas the adapted exoenzyme investment decreases slightly with microsite density. Thus, variation in soil spatial structure has a stronger effect on the evolution of dormancy than on the evolution of exoenzyme production over this parameter range.

The two traits also evolve in opposite directions when mortality parameters vary (Fig. 4E,F). As the death rate of dormant individuals, *d_d_*, increases, dormancy provides less protection, and *ω*\* declines sharply (Fig. 4E). Because cells then become more vulnerable in the bulk soil, selection favors higher exoenzyme investment to ensure minimum survival. Conversely, as the death rate of active individuals in the bulk soil, *d_s_*, increases, dormancy provides relatively more protection, and *ω*\* rises (Fig. 4F). In this case, *φ* * declines because survival in the bulk soil is achieved increasingly through dormancy rather than through exoenzyme production.

## Discussion

We developed a spatially heterogeneous eco-evolutionary model to examine how two microbial traits - entry into dormancy and exoenzyme production - evolve in structured soils and how their evolution feeds back on soil carbon dynamics. The model distinguishes active and dormant microbes in favorable microsites from those in the surrounding bulk soil matrix, and it links microbial growth, exoenzyme-mediated decomposition, and soil carbon stocks. This structure allowed us to ask how selection on microbial life-history traits differs between soil micro-environments and how adaptation at the microbial scale can alter ecosystem-level carbon fluxes.

Both traits have strong, and sometimes counterintuitive, effects on ecological viability and soil carbon storage. Increasing investment in exoenzyme production generally accelerates decomposition and lowers soil carbon stocks. Dormancy has a more non-monotonic effect. At low to moderate levels, increased dormancy can maintain microbial biomass by protecting cells during unfavorable conditions, thereby sustaining decomposition and reducing soil carbon stocks. At higher levels, however, too many individuals remain metabolically inactive, decomposition slows, and soil carbon stocks increase. Thus, the same trait can either amplify or attenuate carbon loss, depending on its value and on the spatial and demographic context in which it evolves.

The evolutionarily adapted value of dormancy investment, *ω*\*, is generated by opposing selective pressures. Within microsites, where growth and resource uptake are possible, dormancy is costly because dormant cells forgo growth. In the bulk soil matrix, where active cells experience high mortality and little or no growth, dormancy is favored because it increases persistence until cells return to suitable microsites. This contrast explains why microbial mobility, microsite density, exoenzyme production, awakening rate, and mortality in the active and dormant states all affect the adaptive level of dormancy. In particular, total movement between microsites and the matrix generally increases the benefit of dormancy, whereas an increase only in movement from the matrix into microsites can reduce the need for dormancy by shortening exposure to the harsh matrix environment.

When dormancy and exoenzyme production jointly evolve, the two traits can either increase together or evolve in opposite directions. Total microbial mobility tends to increase both traits because cells more frequently experience the matrix, where both dormancy and exoenzyme production can improve persistence. In contrast, parameters directly affecting dormancy - such as the cost of entering dormancy, the death rate of dormant individuals, and the death rate of active individuals in the bulk soil - often drive the two traits in opposite directions. This pattern arises because dormancy and exoenzyme production can partly substitute for each other as survival strategies in the matrix. Consequently, the evolution of dormancy can shift the adaptive level of exoenzyme production and thereby alter predictions for soil carbon decomposition.

### Spatial structure and dormancy evolution

Our results connect directly to the classic idea that dormancy can evolve from fine-grained spatial heterogeneity rather than from temporal bet-hedging alone. J. A. J. Metz, Klinkhamer, and Jong 2009 developed such an argument for delayed germination in plants. Their model considers seeds that may germinate in “safe sites,” where seedlings can establish, or outside safe sites, where germination is fatal. Because seeds detect safe sites imperfectly, a strategy that increases germination in safe sites also increases germination outside safe sites. Delayed germination can therefore be favored even in a stable environment, provided that the tradeoff between germination in safe and unsafe sites is sufficiently steep (J. A. J. Metz, Klinkhamer, and Jong 2009).

There is a strong parallel with our model. Soil microsites function as favorable local habitats, similar in spirit to the safe sites of J. A. J. Metz, Klinkhamer, and Jong 2009, whereas the bulk soil matrix is a less favorable environment where active cells face high mortality. In both models, a dormant state allows individuals to avoid the cost of being active in the wrong micro-environment. In both models, the evolutionarily favored strategy is shaped by the balance between exploiting rare favorable sites and avoiding lethal or highly unfavorable conditions outside them. Thus, our model extends the safe-site logic from plant seed banks to microbial life cycles in structured soils.

There are also important differences. In the plant model, the central trait is the probability of germination conditional on location, and delayed germination evolves because imperfect habitat discrimination creates a physiological tradeoff between germinating in safe and unsafe sites. In our model, dormancy evolves because cells move between micro-environments where the same phenotype has different fitness consequences. Dormancy is disfavored in microsites because active growth is possible, but favored in the matrix because active cells die rapidly. In addition, the microbial system includes feedbacks through exoenzyme production, dissolved organic carbon, microbial biomass, and soil carbon pools. These feedbacks mean that adaptive evolution does not merely determine the persistence of a population; it also changes the biogeochemical environment that feeds back on selection. This is a key difference between the present model and the safe-site model of delayed germination.

This comparison also clarifies why soil spatial structure should not be treated as a passive background parameter. In both models, the spatial distribution of favorable sites determines the adaptive value of dormancy. In our case, microsite density and microbial conductance between microsites and the matrix regulate how often microbes experience the contrasting selective regimes. Soil aggregates therefore play a role analogous to evolutionary filters: they shape which microbial strategies persist, and the strategies that persist determine the rate at which carbon is decomposed. In this respect, our results support the view of soil aggregates as “evolutionary incubators” (Rillig, Muller, and Lehmann 2017) and extend that view by showing how the incubator effect interacts with dormancy and exoenzyme-mediated decomposition.

### Dormancy as a social trait

The evolution of microbial dormancy has also been interpreted through the lens of social evolution. Ratcliff et al. 2013 showed that dormancy can have both direct and indirect fitness effects. Dormant microbial cells may survive stresses that kill active cells, but they also spare resources that can be used by active neighbors. In structured populations, these spared resources may be consumed preferentially by clonemates, creating an indirect benefit; in unstructured populations, they may instead be exploited by non-dormant competitors, creating an indirect cost (Ratcliff et al. 2013).

Our model emphasizes a different, but complementary, mechanism. Dormancy evolves here primarily through direct benefits to the focal lineage in the bulk soil matrix: dormant cells avoid the high mortality experienced by active cells in a harsh environment. We do not explicitly model kin structure, assortment among clonemates, or resource sparing as an inclusive-fitness effect. Nevertheless, the contrast with Ratcliff et al. 2013 is informative. In microsites, where cells interact locally and exoenzymes diffuse, microbial traits have strong social consequences. Exoenzyme production is a public good in microsites because non-producing or low-producing strains can benefit from enzymes released by producers. Dormancy could modify this social environment by changing the number of active cells, the amount of resource consumed, and the opportunity for cheaters to exploit extracellular enzymes. Thus, even when dormancy is favored by direct survival benefits, it may reshape the indirect fitness consequences of other cooperative traits.

Twyman and Gardner 2023 provide a broader theoretical framework for this connection by treating dormancy as dispersal through time. They show that kin selection can favor dormancy as a way to reduce competition among relatives, that density-dependent dormancy can generate a “constant non-dormant” principle, and that the relationship between dormancy and the evolution of altruism depends on whether dormancy is density dependent or density independent (Twyman and Gardner 2023). This framework highlights an important dimension not included explicitly in our model: the possibility that dormancy evolves not only because it protects the focal individual from stress, but also because it temporally separates related competitors.

The comparison suggests two interpretations of our results. First, the positive selection for dormancy in the soil matrix can be viewed as a direct stress-tolerance effect, whereas the social-evolutionary models of Ratcliff et al. 2013 and Twyman and Gardner 2023 emphasize indirect effects mediated by resource competition among kin. Second, soil microsites are precisely the type of spatial structure in which the two mechanisms may interact. Limited movement, clonal growth, and local resource diffusion could generate relatedness and assortment within aggregates; at the same time, movement out of aggregates exposes cells to matrix conditions where dormancy has direct survival value. An important extension of our model would therefore be to include explicit genotype structure within microsites and to ask whether dormancy evolves differently when resource sparing and exoenzyme production generate kin-selected benefits or costs.

This social-evolutionary perspective also sharpens the interpretation of joint evolution between dormancy and exoenzyme production. Exoenzyme production is socially costly in microsites because the benefits are shared, but it can be privately beneficial in the matrix if a cell must degrade organic matter near itself to persist. Dormancy changes how long microbes remain in each of these selective contexts. By entering dormancy in the matrix and awakening mainly in microsites, microbes may reduce the time during which exoenzyme production acts as a private survival trait and increase the importance of the microsite public-good phase. This provides a mechanistic explanation for why dormancy can depress the adaptive level of exoenzyme production and thereby protect low-producing strains that would otherwise be counter-selected in the matrix.

### Dormancy and movement through structured soils

Our model also relates to recent theory on the joint evolution of dormancy and dispersal. Cenzer and M’Gonigle 2022 modeled the joint evolution of dormancy and dispersal in spatially autocorrelated landscapes and showed that spatial structure alone can drive the evolution of dormancy. In their simulations, dormancy and dispersal often acted as alternative ways to reduce kin competition; high levels of both traits rarely evolved together, and the favored strategy depended on landscape autocorrelation and dispersal distance (Cenzer and M’Gonigle 2022).

There are substantial similarities with our theory. Both studies show that dormancy can evolve because of spatial structure, not only because of temporal fluctuations. Both studies also emphasize that movement matters: the evolutionary value of dormancy depends on how individuals move relative to favorable and unfavorable parts of the environment. In our model, movement between microsites and the matrix determines how often microbes experience conditions where dormancy is favored. In Cenzer and M’Gonigle 2022, dispersal distance determines how strongly individuals compete with kin and how likely dispersers are to encounter favorable habitat.

The differences are equally important. In the model of Cenzer and M’Gonigle 2022, dormancy and dispersal are alternative mechanisms for avoiding kin competition. In our model, microbial movement and dormancy are not simple substitutes. Movement out of microsites can increase the need for dormancy by exposing cells to the matrix, whereas movement into microsites can reduce the need for dormancy by returning cells to favorable conditions. This asymmetry explains why increasing total mobility can increase dormancy, while increasing only *κ_in_* can decrease it. Moreover, the jointly evolving trait in our model, exoenzyme production, is a resource-acquisition trait that directly affects carbon decomposition. As a result, multi-trait evolution in our model links life-history evolution to ecosystem functioning in a way that is absent from models focused solely on dormancy and dispersal.

These comparisons are conducive to recast the microbial adaptive traits considered here within the Yield-Acquisition-Stress tolerance (Y-A-S) framework (Malik et al. 2020). Exoenzyme production lies primarily on the yield-acquisition axis: increasing *φ* improves access to organic substrates but reduces the carbon available for growth. Dormancy lies on the yield-stress tolerance axis: increasing *ω* improves survival under stress but reduces active growth. In microsites, selection tends to favor yield and to oppose costly investments in dormancy and exoenzyme production, especially when enzymes are shared. In the matrix, selection favors acquisition and stress tolerance because growth is otherwise impossible or mortality is high. The joint evolutionary outcome can therefore move microbial strategies through different regions of the Y-A-S triangle depending on soil structure, mobility, and the severity of matrix conditions.

### Implications for soil carbon and climate change

Because dormancy and exoenzyme production jointly determine microbial activity, their joint evolution may alter predictions of soil carbon feedbacks to climate change. Warming, drying, changes in pore connectivity, altered carbon inputs, and shifts in aggregate structure can all change the relative amount of time microbes spend in microsites versus the bulk soil matrix. These changes are expected to modify selection on dormancy and exoenzyme production, and hence alter decomposition rates. For example, soil drying may increase the selective value of dormancy while reducing microbial movement and extracellular enzyme diffusion. In such cases, dormancy could increase and exoenzyme production could decrease, leading to higher soil carbon stocks than would be predicted from a model with fixed microbial traits. In contrast, soil wetting may reduce dormancy and increase exoenzyme production, potentially accelerating carbon loss.

The direction and magnitude of these effects will depend on the initial ecological state. If most microbes are active, a shift toward dormancy should reduce decomposition. If many microbes are already dormant, a moderate increase in dormancy may instead preserve biomass and maintain decomposition under harsh conditions. Likewise, if exoenzyme production is initially low, small evolutionary changes in *φ* can have disproportionately large effects on dormancy selection and carbon stocks. These nonlinearities caution against representing dormancy as a fixed inactive fraction or exoenzyme production as a fixed decomposition parameter in soil carbon models.

## Conclusion

Our results therefore suggest that models neglecting multi-trait microbial eco-evolutionary feedbacks may either underpredict or overpredict climate-induced soil carbon losses. The sign of the error depends on soil structure, microbial mobility, dormancy costs, awakening rates, and the joint evolutionary response of exoenzyme production and dormancy. Empirically, this points to several priorities: estimating the active-to-dormant ratio across microhabitats; measuring rates of movement into and out of soil aggregates; quantifying how dormancy and exoenzyme production covary across genotypes and environments; and determining how kin structure within microsites affects resource sharing and public-good dynamics.

More generally, our study unites previous lines of theory (spatial heterogeneity and dormancy evolution, dormancy as a social trait, and eco-evolutionary feedbacks mediated by microbial decomposition), a common theme being that dormancy evolves in response to heterogeneity in the fitness value of activity. In soils, that heterogeneity is temporal, spatial, social, and biogeochemical. By coupling microbial adaptive dynamics to carbon cycling, our model shows that the evolution of dormancy is not merely a question of microbial life-history adaptation. Through direct effects on microbial activity and interactions and indirect effects on the evolutionary adaptation of exoenzyme production, it is also potentially a major regulator of soil-atmosphere carbon feedbacks.

## Acknowledgments

A.C.-L. is supported by the French Ministry of Agriculture for her doctoral research. R.F. is supported by the Biology Integration Institute EMERGE, U.S. National Science Foundation (NSF) Biology Integration Institutes Program, award # 2022070, and by the NSF Growing Convergence Research, award 2121155. We thank Elsa Abs, Antonin Affholder and Scott Saleska for discussion.

## Statement of Authorship

A.C.-L. and R.F. conceptualized the study. A.C.-L. developed the model, did the math and coding, and generated the results. A.C.-L. and R.F. interpreted the results. A.C.-L. wrote the first draft of the manuscript, which was revised by R.F. Both authors finalized the manuscript.

## Supplementary Figures

**Figure S1.**
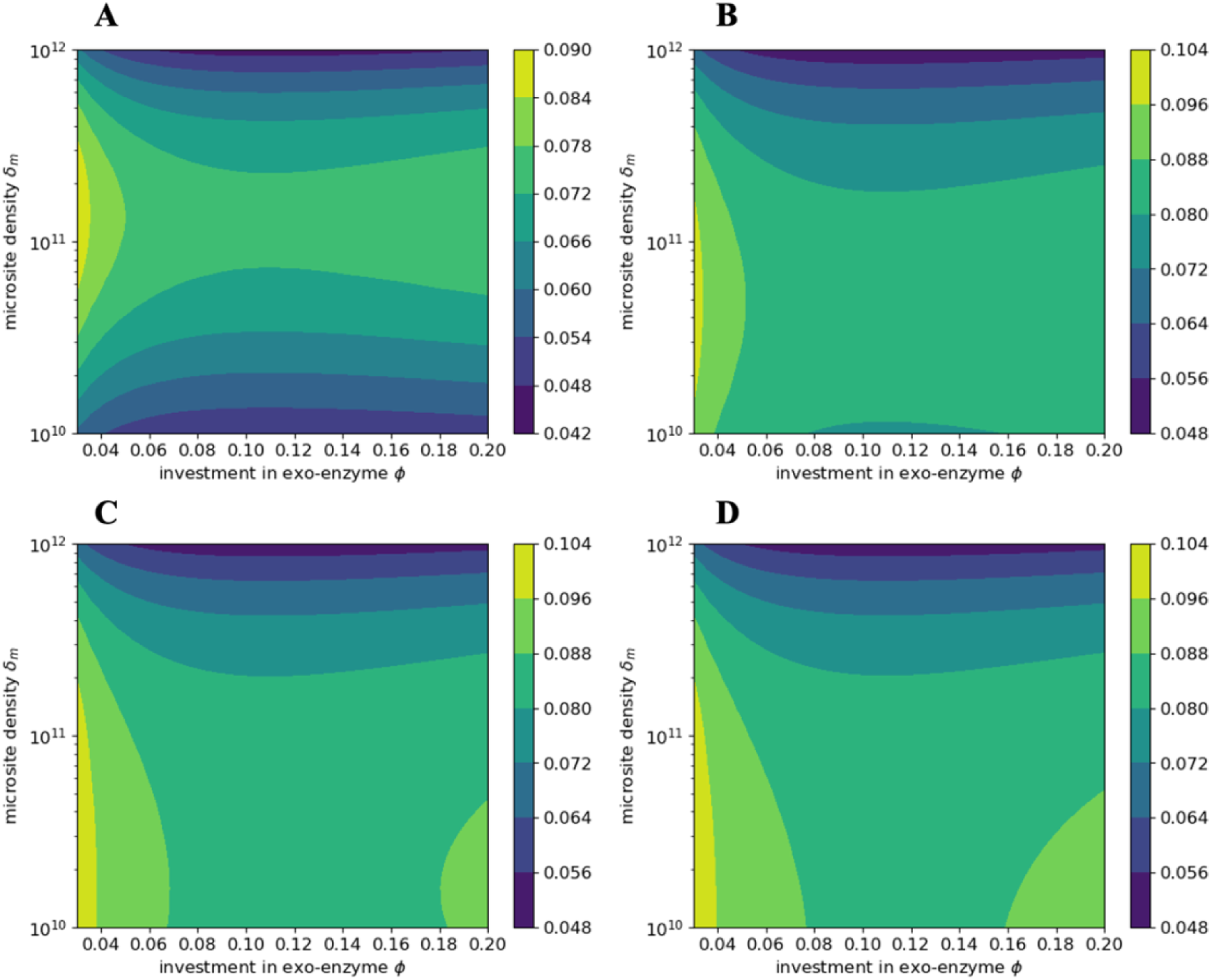
Evolutionarily adapted dormancy entry. Adapted investment in dormancy, ω*, as a function of exoenzyme investment, *φ* and microsite density *δ_m_* for (A) *d_d_* = 10^−6^; (B) *d_d_* = 10^−7^; (C) *d_d_* = 10^−8^ and (D) *d_d_* = 10^−9^.

**Figure S2.**
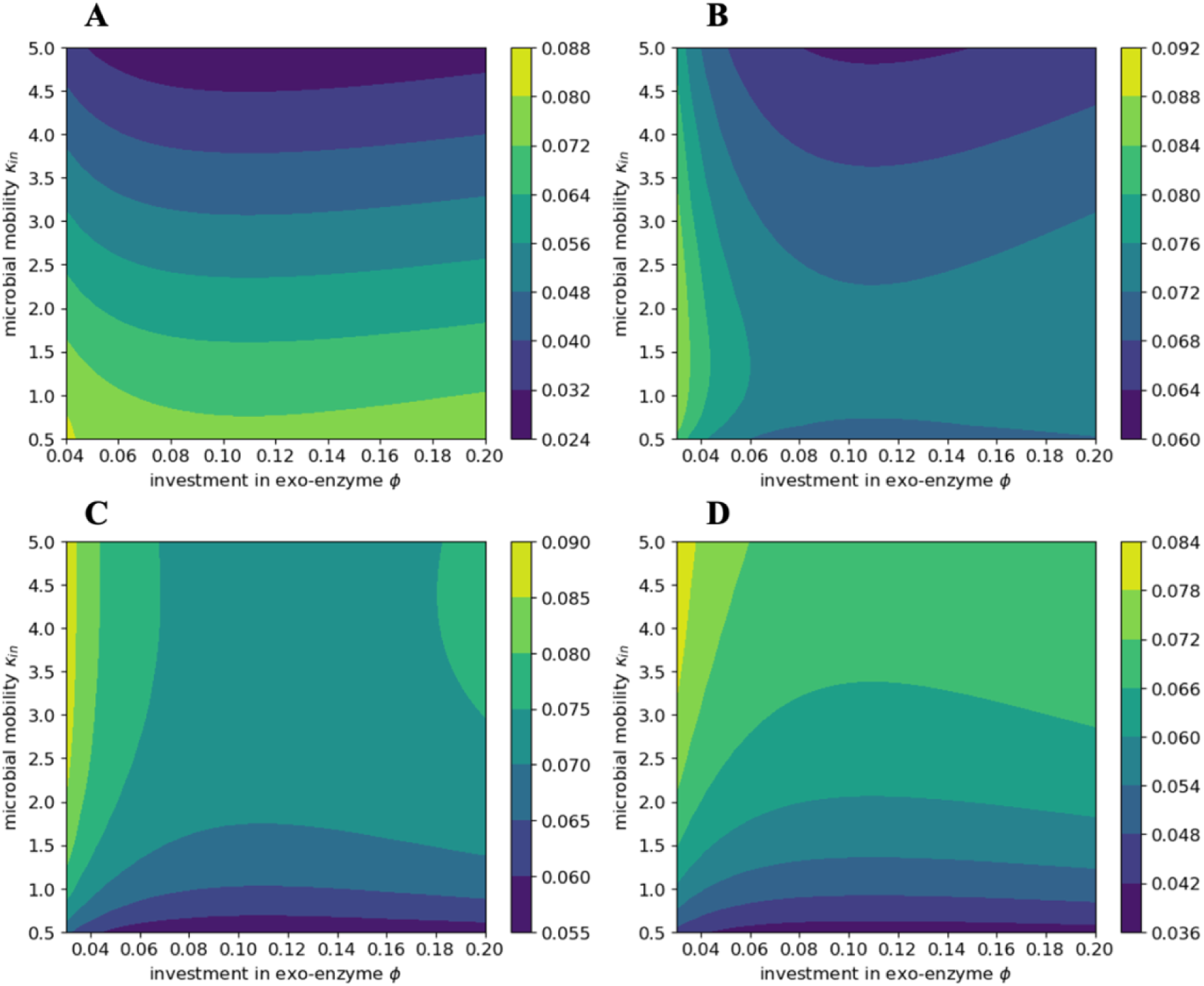
Evolutionarily adapted dormancy entry. Adapted investment in dormancy, ω*, as a function of exoenzyme investment, *φ* and microbial mobility *κ_in_* for: (A) *δ_m_* = 3 × 10^11^m^-3^; (B) *δ_m_* = 10^11^m^-3^; (C) *δ_m_* = 3 × 10^10^m^-3^ and (D) *δ_m_* = 10^10^m^-3^.

**Figure S3.**
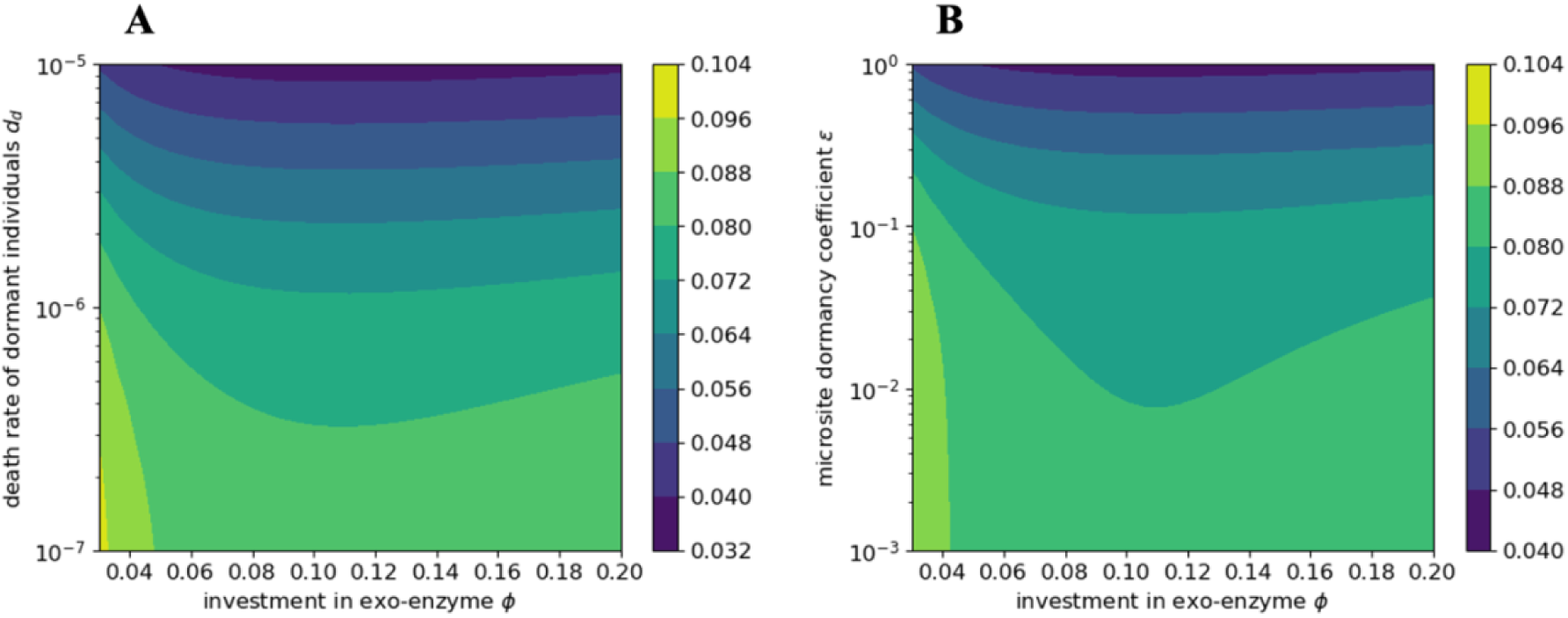
Evolutionarily adapted dormancy entry. Adapted investment in dormancy, ω*, as a function of exoenzyme investment, *φ* and (A) death rate of dormant individuals *d_d_*; (B) microsite dormancy coefficient *ε*.

## Notes

### Competing Interest Statement

The authors have declared no competing interest.

